# Two point mutations in the Hantaan virus glycoproteins afford the generation of a highly infectious recombinant vesicular stomatitis virus vector

**DOI:** 10.1101/356055

**Authors:** Megan M. Slough, Kartik Chandran, Rohit K. Jangra

**Author notes:** Corresponding authors. (K.C.); (R.K.J.).

## Abstract

Rodent-to-human transmission of hantaviruses is associated with severe disease. Currently, no FDA-approved, specific antivirals or vaccines are available, and the requirement for high biocontainment (BSL3) laboratories limits hantavirus research. To study hantavirus entry in a BSL-2 laboratory, we set out to generate replication-competent, recombinant vesicular stomatitis viruses (rVSVs) bearing the Gn/Gc entry glycoproteins. As previously reported, rVSVs bearing New World hantavirus Gn/Gc were readily rescued from cDNAs, but their counterparts bearing Gn/Gc from the Old World hantavirus, Hantaan virus (HTNV), were refractory to rescue and only grew to low titers. However, serial passage of the rescued rVSV-HTNV Gn/Gc virus markedly increased its infectivity and capacity for cell-to-cell spread. This gain in viral fitness was associated with the acquisition of two point mutations; I532K in the cytoplasmic tail of Gn, and S1094L in the membrane-proximal stem of Gc. Follow-up experiments with rVSVs and single-cycle VSV pseudotypes confirmed these results. Mechanistic studies revealed that both mutations were determinative and contributed to viral infectivity in a synergistic manner. Our findings indicate that the primary mode of action of these mutations is to relocalize HTNV Gn/Gc from the Golgi complex to the cell surface, thereby affording significantly enhanced Gn/Gc incorporation into budding VSV particles. Our results suggest that enhancements in cell-surface expression of hantaviral glycoprotein(s) through incorporation of cognate mutations could afford the generation of rVSVs that are otherwise challenging to rescue. The robust replication-competent rVSV-HTNV Gn/Gc reported herein may also have utility as a vaccine.

**Importance:** Human hantavirus infections cause pulmonary syndrome in the Americas and hemorrhagic fever with renal syndrome (HFRS) in Eurasia. No FDA-approved vaccines and therapeutics exist for these deadly viruses, and their development is limited by the requirement for high biocontainment. In this study, we identified and characterized key amino acid changes in the surface glycoproteins of HFRS-causing Hantaan virus that enhance their incorporation into recombinant vesicular stomatitis virus (rVSV) particles. The replication-competent rVSV genetically encoding Hantaan virus glycoproteins described in this work provides a powerful and facile system to study hantavirus entry under lower biocontainment and may have utility as a hantavirus vaccine.

## Introduction

Rodent-borne hantaviruses (family *Hantaviridae* of segmented negative-strand RNA viruses) cause hemorrhagic fever with renal syndrome (HFRS) in Eurasia and hantavirus pulmonary syndrome (HPS) in the Americas (1). Globally, more than 150,000 cases of hantavirus disease occur per year. Human population growth, accelerating climate change, and habitat loss are predicted to increase the size and severity of hantavirus disease outbreaks (2–6). Although inactivated viral vaccines are in use in Asia for HFRS-causing Seoul (SEOV) and Hantaan (HTNV) viruses, their protective efficacy is moderate at best (7–9), and no FDA-approved hantavirus vaccines or antivirals are available. The development of hantavirus countermeasures is hampered by our limited understanding of the molecular mechanisms of viral replication and disease pathogenesis, the lack of tools available to investigate these mechanisms, and the need to perform hantavirus research under high biocontainment.

The development of surrogate viral systems (10, 11) that recapitulate cell entry and infection under BSL-2 containment (or lower) has greatly accelerated both basic mechanistic investigations of virulent emerging viruses and the discovery and development of vaccines and therapeutics to target them (12–15). Several such systems have been described for hantaviruses, whose glycoproteins Gn and Gc (hereafter, Gn/Gc) are necessary and sufficient for viral entry (16). When expressed in cells with or without the nucleoprotein N, Gn/Gc were shown to self-assemble to produce virus-like particles (VLPs) with utility for studies of viral glycoprotein maturation and assembly (17–19) and as potential vaccine vectors (18). Single-cycle gammaretroviral and lentiviral vectors bearing HTNV or ANDV Gn/Gc have been employed for viral entry and antibody neutralization studies, and as candidate vectors for vaccination and gene therapy (20–24). Consistent with the flexibility of heterologous protein incorporation in the budding virions of vesicular stomatitis virus (VSV), multiple groups have also developed VSV-based single-cycle pseudovirions for both HFRS-causing hantaviruses [HTNV (16, 25–27), Puumala virus (PUUV) (26, 28) and SEOV (16)] and HPS-causing hantaviruses [ANDV (29) and Sin Nombre virus (SNV) (27)].

Although single-cycle pseudovirions have advanced our understanding of hantavirus Gn/Gc assembly, viral entry, and antiviral immune responses, they are labor-intensive to generate in high yield. By contrast, self-replicating, recombinant VSVs (hereafter, rVSVs), whose genomes have been modified to carry the hantavirus M gene (encoding Gn/Gc) in place of the VSV glycoprotein (G) gene, are relatively easy to produce in quantity, readily amenable to forward-genetic and small-molecule screens, and are unique among surrogate systems in affording forward-genetic selections to identify escape mutants against neutralizing antibodies and small-molecule entry inhibitors (30–33). Brown *et al.* (34), were the first to generate a rVSV bearing ANDV Gn/Gc and showed that it could protect Syrian hamsters against lethal ANDV challenge when administered as a vaccine (34, 35). Similar viruses have been used to identify host factors required for ANDV entry (27, 36). To expand the pool of such rVSVs for hantavirus research, we previously rescued a rVSV bearing SNV Gn/Gc from cDNA (36). However, rVSVs bearing Gn/Gc from the HFRS-causing Hantaan virus (HTNV) proved challenging to rescue and yielded only a slowly replicating virus.

Here, we show that serial passage of the initial rVSV-HTNV Gn/Gc stock in cell culture afforded the generation of a variant with enhanced replicative fitness suitable for viral entry studies. We mapped this gain in viral fitness to the acquisition of two point mutations: I532K in the cytoplasmic tail of Gn, and S1094L in the stem region of Gc. Mechanistic studies revealed that these mutations enhance rVSV infectivity by relocalizing HTNV Gn/Gc from the Golgi complex to the cell surface, thereby augmenting Gn/Gc incorporation into budding VSV particles. Our results suggest that selection- and protein engineering-based approaches to boost the cell surface expression of entry glycoproteins from other hantaviruses, or even more divergent bunyaviruses, could enable the generation of rVSVs that are otherwise refractory to rescue and/or replicate only poorly. The rVSV-HTNV Gn/Gc vector described herein may have utility as an HTNV vaccine.

## Results

### Two point mutations in the Gn/Gc complex enhance spread and replication of rVSV-HTNV Gn/Gc

Hantaviruses are classified as biosafety level-3 (BSL-3) agents. To study hantavirus entry and infection in a BSL-2 setting, we attempted to generate a replication-competent, recombinant vesicular stomatitis virus (rVSV) expressing the Gn/Gc glycoproteins of Hantaan virus (HTNV), a prototypic HFRS-causing hantavirus. Multiplication and spread of the early-passage rVSV bearing HTNV Gn/Gc was poor but improved dramatically following three serial passages in Vero cells. Analysis of the selected viral population identified two amino acid changes in Gn/Gc, one located in the cytoplasmic tail of Gn (I532K) and the other in the membrane-proximal stem of the Gc ectodomain (S1094L) (**Fig. 1A**). To determine if either or both mutations could account for enhanced viral multiplication, we attempted to rescue rVSV-HTNV Gn/Gc viruses from cDNAs incorporating each mutation separately and together. We successfully recovered both single- and double-mutant viruses, but not the rVSV bearing WT HTNV Gn/Gc. Growth curves revealed that the Gn/Gc double-mutant virus (I532K/S1094L) multiplied and spread more rapidly than either single-mutant virus (**Fig. 1B** and **C**).

**Fig. 1.**
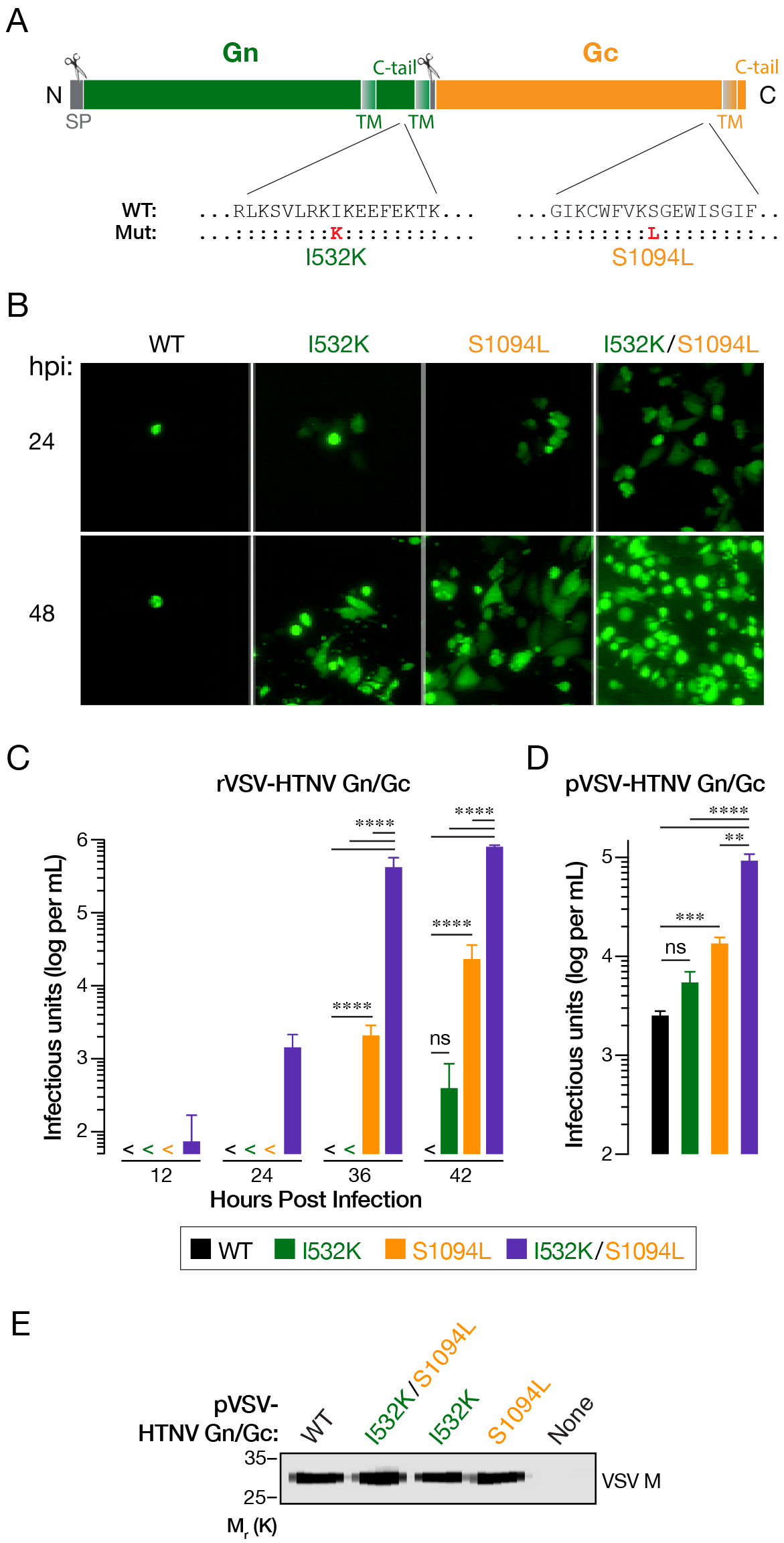
Two point mutations (I532K & S1094L) in the Gn/Gc enhance rVSV-HTNV Gn/Gc spread and replication. (A) Schematic representation of the HTNV Gn/Gc. Location of the point mutations acquired after serial passaging of rVSV expressing HTNV Gn/Gc. (B-C) Growth of WT or mutant rVSV-HTNV Gn/Gc. Supernatants from 293FT cells co-transfected with plasmids encoding rVSV genomes expressing eGFP bearing WT, I532K, S1094L or I532K/S1094L versions of HTNV Gn/Gc with helper plasmids, were used to infect Vero cells. (B) Representative images of eGFP expression in Vero cells at indicated times post infection. (C) Supernatants collected from infected Vero cells at indicated times post-infection were titered on naive Vero cells. Data from two independent experiments (n = 4) are represented as log infectious units (IU) per mL (mean ± SD). “>” indicate virus titers that were below the limit of detection (50 IU per mL). Groups were compared by two-way ANOVA with Tukey’s correction for multiple comparisons. ns (not significant), *P* > 0.05; ****, *P* < 0.0001. (D) Production of single VSV pseudotypes (pVSV). 293T cells expressing WT, I532K, S1094L or I532K/S1094L forms of HTNV Gn/Gc *in trans*, were infected with VSV-eGFP-AG (VSV expressing eGFP and carrying VSV G glycoprotein on its surface, but lacking the G gene) 48 h later. Following extensive washing to remove VSV G-carrying residual viruses, supernatants were collected at 48 h post-infection and infectious titers were measured on Vero cells. Mean ± SEM from 4 independent experiments (n = 8) are shown here. Background VSV pseudotype production from empty vector-transfected cells was below the limit of detection (100 IU per mL). Groups were compared by one-way ANOVA with Tukey’s correction for multiple comparisons. ns, *P* > 0.05; **, *P* < 0.01; ***, *P* < 0.001; ****, *P* < 0.0001. (E) HTNV Gn/Gc mutations do not affect overall VSV particle production. Equivalent amounts (by volume) of pelleted VSV pseudotypes from panel D were analyzed by VSV M-specific immunoblotting. A representative blot from 3 independent experiments is shown.

To confirm the determinative role of each Gn/Gc mutation and exclude potentially confounding effects from mutations elsewhere in the viral genome, we generated and analyzed single-cycle VSV pseudotypes (pVSVs) bearing WT and mutant Gn/Gc proteins. Consistent with our findings with the rVSVs, pVSVs bearing the HTNV Gn/Gc (I532K/S1094L) double-mutant displayed a higher specific (per-particle) infectivity in Vero cells than those bearing either single-mutant or WT Gn/Gc (**Fig. 1D**), after normalization of each viral preparation for particle number (**Fig. 1E**). Further, each single-mutant was more infectious than WT. Together, these findings indicate that mutations I532K and S1094 in HTNV Gn and Gc, respectively, make individual contributions to VSV-HTNV Gn/Gc infection, but confer a synergistic enhancement in infectivity when present in the same viral particles.

### The I532K/S1094L double mutation modestly enhances Gn/Gc production

To uncover the mechanism by which the mutations I532K and S1094L enhance VSV-HTNV Gn/Gc infectivity, we first examined the effects of these mutations on Gn/Gc expression. We transfected 293T cells with plasmids encoding WT or mutant Gn/Gc and used a cell-based ELISA to determine the relative Gn/Gc levels (see **Materials and Methods** for details). Cells were permeabilized to render Gn/Gc in all subcellular compartments accessible to immunodetection by the conformation-sensitive, HTNV Gc-specific monoclonal antibody 3G1 (37). At 48 h post-transfection, the I532K/S1094L double mutation modestly elevated Gn/Gc expression in comparison to WT, although I532K had little or no effect, and S1094L modestly depressed Gn/Gc expression (**Fig. 2**). However, none of these changes from WT were statistically significant. These findings suggest that an increase in the steady state levels of Gn/Gc is not the primary mechanism by which the I532K/S1094L mutations enhance rVSV-HTNV Gn/Gc infectivity.

**Fig. 2.**
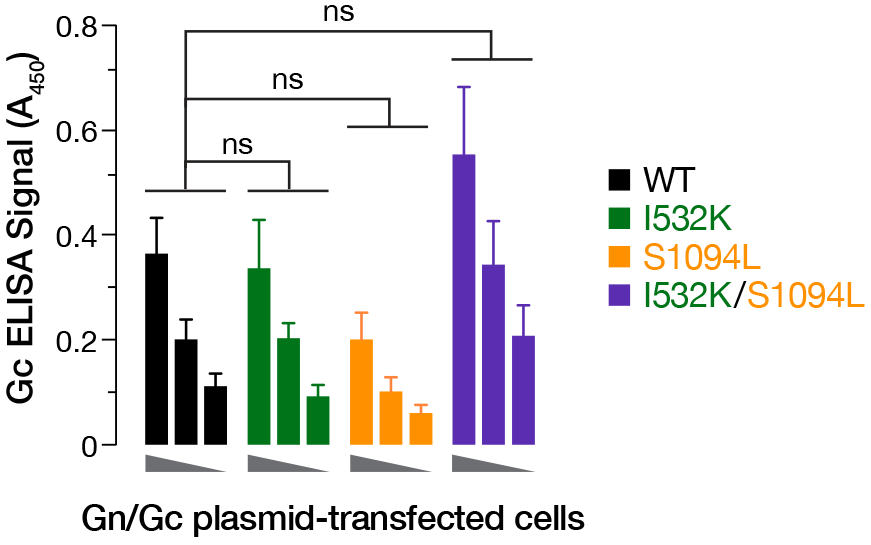
The HTNV Gn/Gc double mutations modestly enhances Gc production. 293T cells, transfected with plasmids expressing empty vector or wild type, I532K, S1094L, or I532K/S1094L forms of HTNV Gn/Gc, were fixed at 48 h post-transfection. Following permeabilization, cells were tested for HTNV total Gc expression using an in-cell ELISA. To ensure linearity of the ELISA, serial 2-fold dilutions of the transfected cells were made by mixing them with untransfected cells. Data were graphed after background ELISA signal from empty vector-transfected 293T cells was subtracted. Data from 3 independent experiments (n = 5-6) are shown as mean ± SEM. Groups were compared by one-way ANOVA with Tukey’s correction for multiple comparisons. ns, *P* > 0.05.

### The I532K/S1094L mutations together enhance localization of Gn/Gc at the plasma membrane

Both VSV and HTNV acquire their surface glycoproteins in the secretory pathway during viral budding; however, they do so in distinct cellular compartments. Specifically, VSV particles bud at the plasma membrane, whereas HTNV particles, like those of many hantaviruses and other bunyaviruses, are reported to bud into the Golgi apparatus. In keeping with these observations, VSV G and HTNV Gn/Gc (**Figs 3–4**) localize primarily to the plasma membrane and endoplasmic reticulum (ER)/Golgi apparatus, respectively (38–40). We postulated that this mismatch in the subcellular sites of viral budding and glycoprotein localization may account for the poor growth of rVSV-HTNV Gn/Gc, and that the I532K/S1094L mutations might ameliorate this mismatch. Accordingly, we examined the subcellular distribution of WT and mutant Gn and Gc in transfected U2OS cells by immunofluorescence (IF) microscopy. Gn and Gc colocalization was essentially complete for all variants (**Fig. 3**), showing that the mutations do not alter relative Gn and Gc distribution in cells. We further noted a predominantly perinuclear Gn/Gc staining for all variants that colocalized with GM130, a marker for the Golgi apparatus (**Fig. 4**), indicating that the mutants substantially retain Golgi localization. Interestingly, however, some cells expressing the double mutant also displayed marked Gn/Gc staining in the cell periphery (yellow arrows in **Figs. 3** and **4**), suggesting that a subset of these molecules do localize to the plasma membrane.

**Fig. 3.**
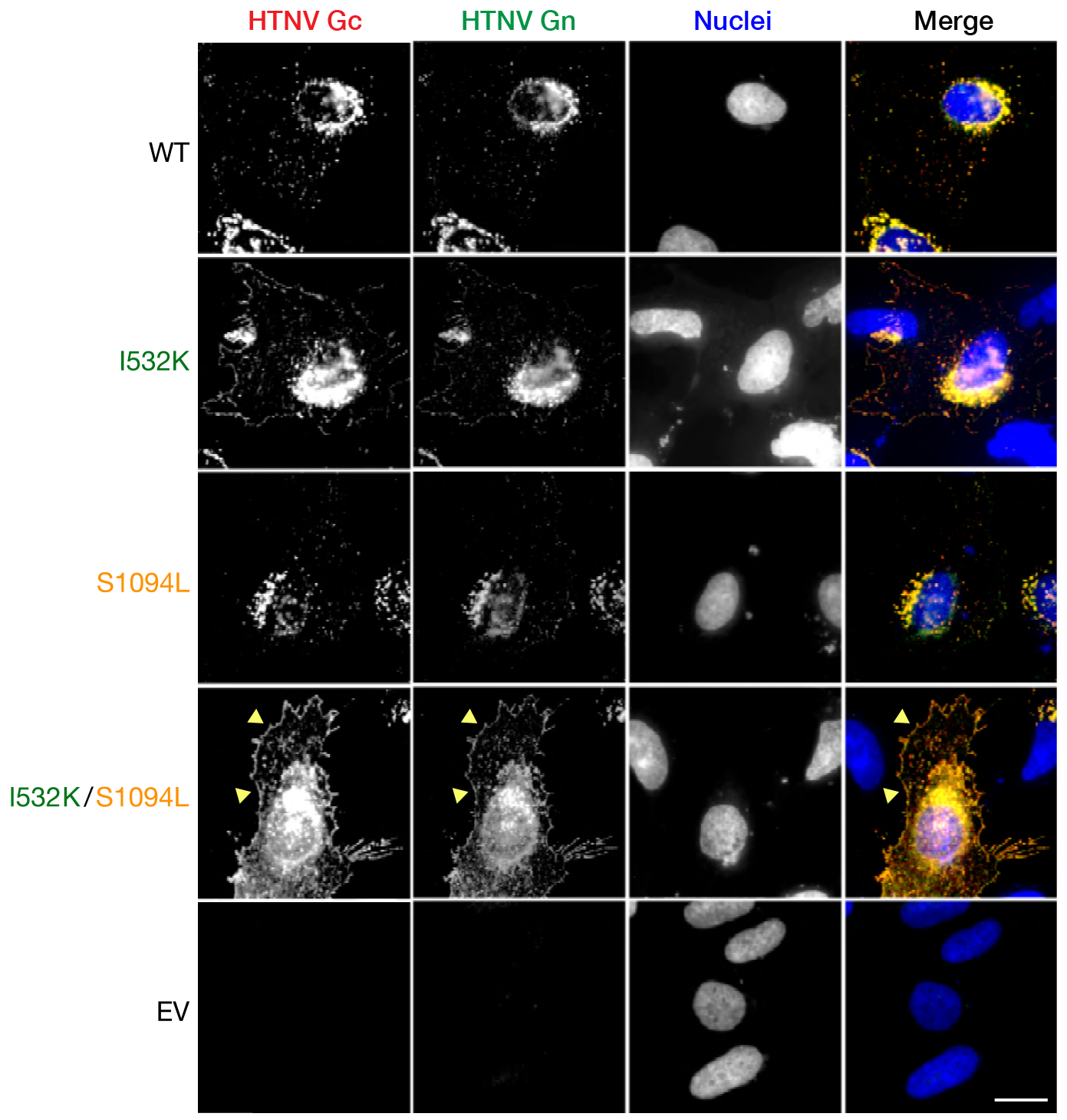
The HTNV Gn/Gc mutations do not alter glycoprotein co-localization. Human osteosarcoma U2OS cells, transfected with plasmids expressing wild type, I532K, S1094L, or I532K/S1094L variants of HTNV Gn/Gc, were fixed at 24 h post-transfection, permeabilized and co-immunostained with HTNV Gn- and Gc-specific antibodies (see Materials and Methods for details). EV, empty vector. Representative images for each of the HTNV variants from an experiment representative of at least 3 experiments are shown. Scale bar, 20 μm.

**Fig. 4.**
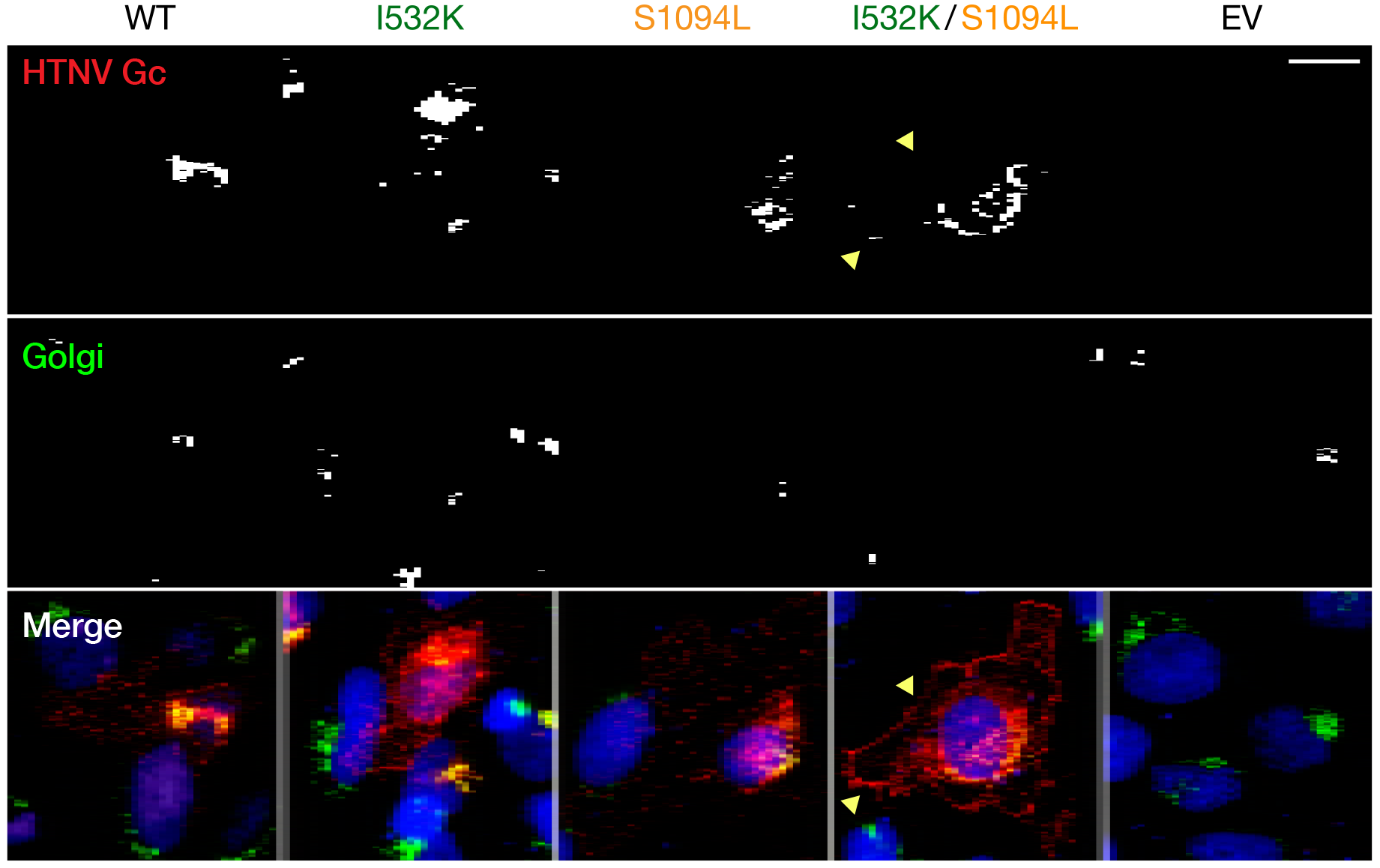
Trafficking of HTNV Gn/Gc to the Golgi apparatus is unaffected by I532K & S1094L mutations. (A) Human osteosarcoma U2OS cells, transfected with plasmids expressing wild type, I532K, S1094L, or I532K/S1094L forms of HTNV Gn/Gc, were fixed at 24 h post-transfection, permeabilized, and co-stained with HTNV Gc specific (3G1) and Golgi apparatus-specific (GM130) antibodies. EV, empty vector. Representative images from an experiment, out of at least 3 independent experiments, are shown. Scale bar, 20 μm.

To directly examine this possibility, we immunostained U2OS cells transfected with plasmids expressing WT or mutant Gn/Gc to visualize the cell surface expression of these glycoproteins. The Gn mutation alone enhanced the cell-surface expression of both Gn and Gc, whereas the Gc mutation alone had little or no effect. Interestingly, the double-mutant afforded an even higher level of cell-surface Gn/Gc expression, indicating that the Gc mutation can act in concert with its Gn counterpart to drive relocalization of Gn/Gc to the plasma membrane (**Fig. 5A**). Similar results were obtained with primary human umbilical vein endothelial cells (HUVEC) transfected with HTNV Gn/Gc expression plasmids (**Fig. 5B**). Quantitation of cell-surface Gc expression in plasmid-transfected U2OS cells by flow cytometry (**Fig. 5C**), and 293T cells by on-cell ELISA (**Fig. 5D**), further corroborated the synergistic enhancement of Gn/Gc cell-surface expression conferred by the I532K and S1094L mutations.

**Fig. 5.**
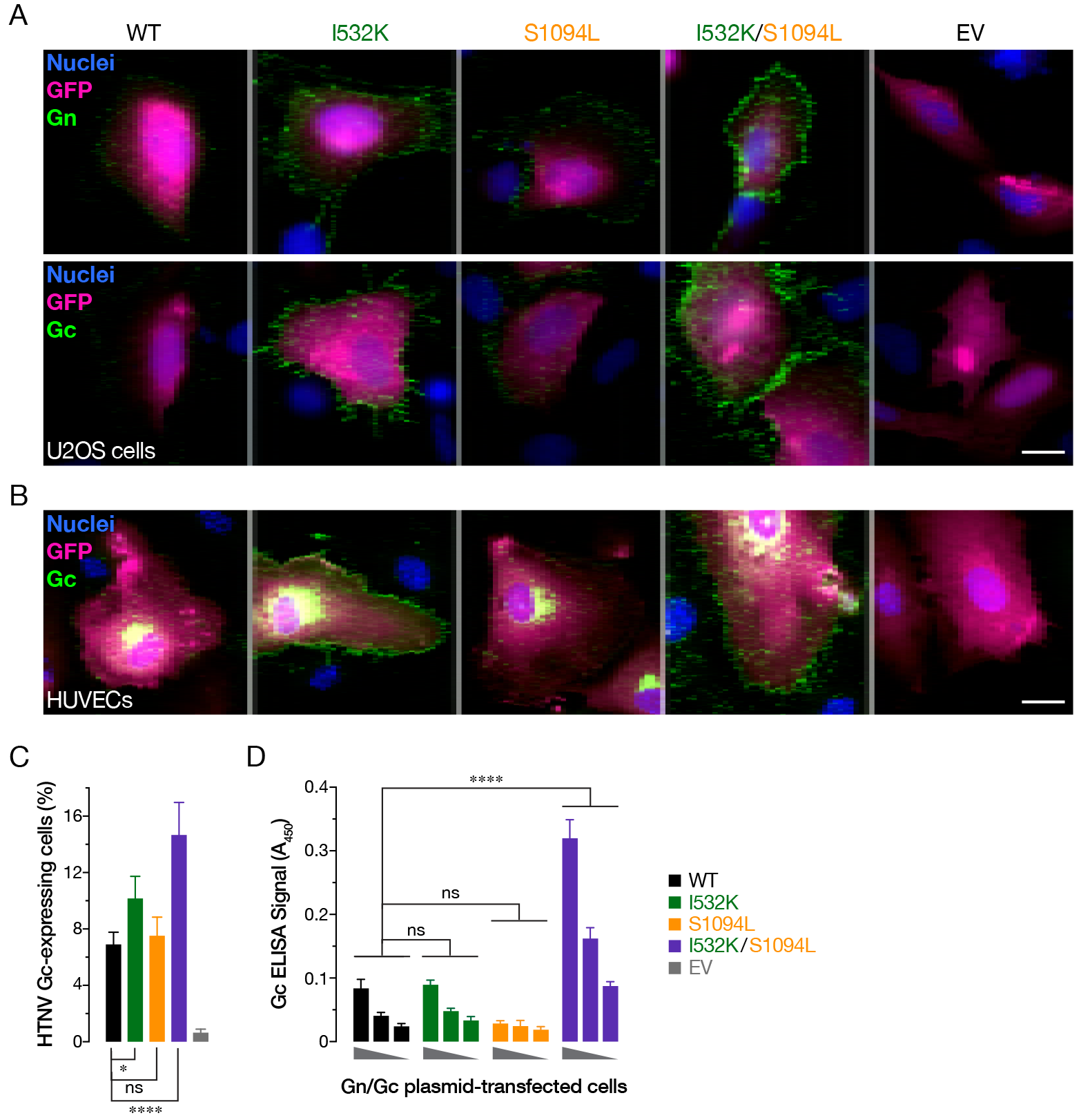
The HTNV Gn/Gc I532K & S1094L mutations together enhance plasma membrane localization of Gn/Gc. (A) U2OS cells, co-transfected with plasmids expressing eGFP and wild type, I532K, S1094L or I532K/S1094L forms of HTNV Gn/Gc, were stained for cell surface expression of HTNV Gn or Gc at 48 h post-transfection. (B) Primary human endothelial cells (HUVECs), nucleofected with plasmids encoding eGFP and wild type, I532K, S1094L, or I532K/S1094L versions of HTNV Gn/Gc, were fixed 72 h later, permeabilized, and stained with HTNV Gc-specific antibody. Representative images from a single experiment, illustrating at least 3 independent experiments, are shown for each panel A and B. EV, empty vector. Scale bars, 20 μm. (C) U2OS cells, transfected as described in panel A, were immunostained for cell surface expression of HTNV Gc and analyzed using flow cytometry. Data from 3 independent experiments are shown as mean ± SD. Groups were compared by one-way ANOVA with Tukey’s correction for multiple comparisons. ns, *P* > 0.05; *, *P* < 0.05; ****, *P* < 0.0001. (D) 293T cells, transfected with plasmids expressing variants of HTNV Gn/Gc, were stained, at 48 h post-transfection, for cell surface expression of HTNV Gc, and detected by on-cell ELISA using Gc-specific mAb 3G1 (Mean ± SEM, n = 5-6 from 3 independent experiments). Groups were compared by two-way ANOVA with Tukey’s correction for multiple comparisons. ns, *P* > 0.05; ****, *P* < 0.0001.

### HTNV-Gn/Gc mutations collectively increase viral glycoprotein incorporation into VSV virions

Because VSV virions are known to acquire heterologous membrane proteins during viral budding at the host-cell plasma membrane, we reasoned that enhanced cell-surface–expression of mutant HTNV Gn/Gc might increase incorporation of the latter into VSV particles. To test this hypothesis, we examined single-cycle VSV pseudotypes (pVSVs) bearing WT and mutant Gn/Gc proteins for HTNV Gn/Gc incorporation by HTNV Gc-specific ELISA, after normalizing viral particle content by VSV matrix protein M-specific immunoblotting (**Fig. 6A**). Concordant with their effects on the cell-surface expression level of each glycoprotein, the Gn mutation alone enhanced incorporation of Gn/Gc into viral particles, whereas the Gc mutation did not, and combination of both mutations afforded a further synergistic increase in Gn/Gc incorporation (**Fig. 6B**). Taken together, these findings strongly suggest that relocalization of HTNV Gn/Gc from the Golgi complex to the plasma membrane induced by the I532K/S1094L mutations enhances rVSV-HTNV Gn/Gc infectivity by increasing viral glycoprotein incorporation into virus particles.

**Fig. 6.**
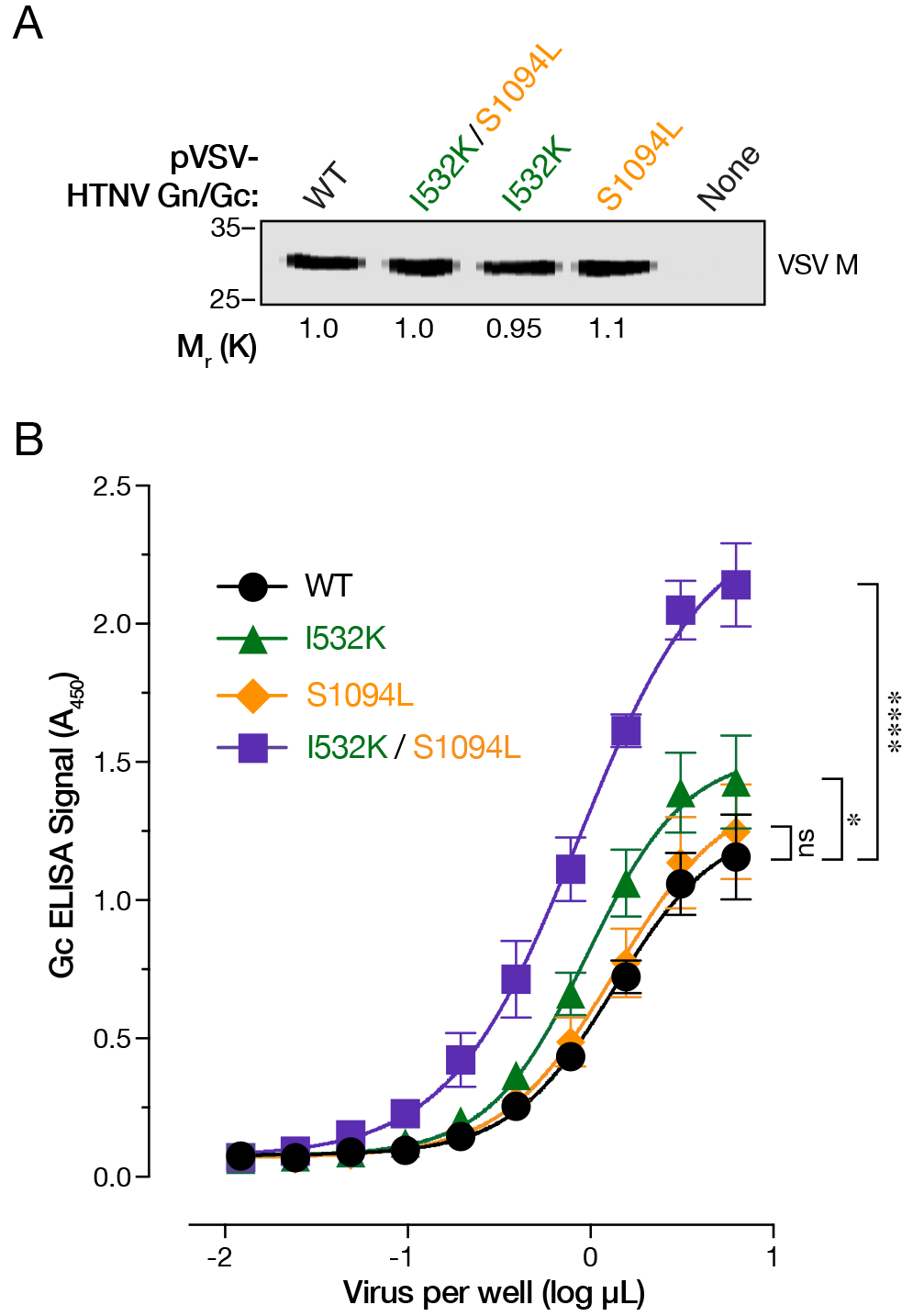
I532K & S1094L mutations collectively increased HTNV Gn/Gc incorporation into VSV virions. (A) Normalization of virus particles carrying HTNV Gn/Gc variants by immunoblotting of VSV matrix (M) protein. Numbers below the blot indicate relative amount of VSV M protein detected as compared to the WT. Representative blot of at least 3 independent experiments is shown here. (B) Serial 2-fold dilutions of normalized pVSV particles as ascertained in panel A, were captured on an ELISA plate and subjected to an HTNV Gc-specific ELISA (Mean ± SEM, n = 8 from 4 independent assays performed on two independent virus preparations). Groups were compared by two-way ANOVA with Tukey’s correction for multiple comparisons. ns, *P* > 0.05; ****, *P* < 0.0001.

## Discussion

Retroviral and vesiculoviral pseudotypes carrying heterologous viral glycoproteins have greatly enhanced our understanding of viral glycoprotein maturation and virus assembly (20, 21), helped delineate roles of host factors in viral entry and other virus-host interactions (36, 41–43), assisted decipher mechanisms of immune response and correlates of protection (34, 44), and have successfully been used to isolate and characterize neutralizing antibodies (31, 45) and developed as vaccines (15, 34). Notwithstanding the remarkable ability of these virions to package heterologous glycoproteins that localize to the plasma membrane, not all viral entry glycoproteins are amenable to efficient pseudotyped virus production. Here, we combined the remarkable ability of rVSV to undergo mutations, akin to other RNA viruses, with forward genetic analyses to identify and characterize the role of two point mutations, one each in the HTNV Gn (I532K) and Gc (S1094L), that greatly enhance infectivity.

Like most members of the order *Bunyavirales*, HFRS-causing HTNV has been shown to bud at the Golgi cisternae (46, 47), with undetectable (48) or very low (16, 48, 49) amounts of Gn/Gc observed at the surface of cells expressing HTNV or another Old World hantavirus SEOV Gn/Gc. We hypothesized that I532K/S1094L mutations facilitate rVSV rescue by altering Gn/Gc expression and/or localization. Both of these mutations alone or together did not significantly affect total protein production (**Fig. 2**) or colocalization of Gn and Gc (**Fig. 3**). Although the majority of the single or double mutant Gn/Gc proteins were still localized to the Golgi complex (**Fig. 3**), as were the WT proteins, the Gn mutation alone (I532K) or together with the Gc mutation (I532K/S1094L) showed significantly elevated cell-surface expression as seen by immunofluorescence (**Figs. 3–4, 5A–B**), flow cytometry (**Fig. 5C**) and on-cell ELISA (**Fig. 5C**). As observed previously (16, 49), we also see some WT HTNV Gn/Gc protein expression on the cell surface (**Figs. 3–5**). However, the I532K/S1094L mutations consistently enhanced cell surface expression, by 3- to 4-fold as compared to the WT, in multiple human cell lines (U2OS and 293T), as well as primary cells (HUVECs), at multiple times post-transfection, suggesting that this phenotype is not limited to a particular cell type or time point (**Figs. 3–5**). Importantly, enriched cell-surface expression correlated well with the levels of HTNV Gn/Gc incorporated in the vesiculoviral pseudovirions (**Fig. 6**), strongly indicating that relocalization of Gn/Gc from Golgi complex to the cell surface is the major mechanism by which these mutations enhance rVSV-HTNV Gn/Gc infectivity. Moreover, the rVSV-HTNV Gn/Gc resembled the authentic HTNV (36) with respect to dependence on the sterol regulatory element-binding protein (SREBP) pathway and cholesterol requirements for entry and infection (27, 36) underscoring its utility for studying hantavirus entry.

Some bunyaviral glycoproteins, including those of hantaviruses, are expressed on the cell surface of virus-infected as well as glycoprotein cDNA-transfected cells (50–54). Consistent with the localization of readily detectable Gn/Gc on the cell surface of HPS-causing viruses (53, 55), transmission electron microscopic studies show evidence of plasma-membrane assembly of some New World hantaviruses such as SNV and Black Creek Canal virus (BCCV) (55, 56). Moreover, ANDV or SNV Gn and Gc can replace each other without affecting their normal trafficking (57). Interestingly, Gn of the HTNV, SEOV & another HRFS-causing Dobrava-Belgrade virus (DOBV) carries isoleucine at position 532 (PUUV is an exception), but it is a valine in that of the New World hantaviruses (**Fig. 7**). Congruent with these differences in cellular localization of their glycoproteins and virion budding sites, rescue of replication- and propagation-competent rVSVs carrying Gn/Gc from ANDV (34, 36) or SNV (36) was relatively easier than those carrying HTNV Gn/Gc.

**Fig. 7.**
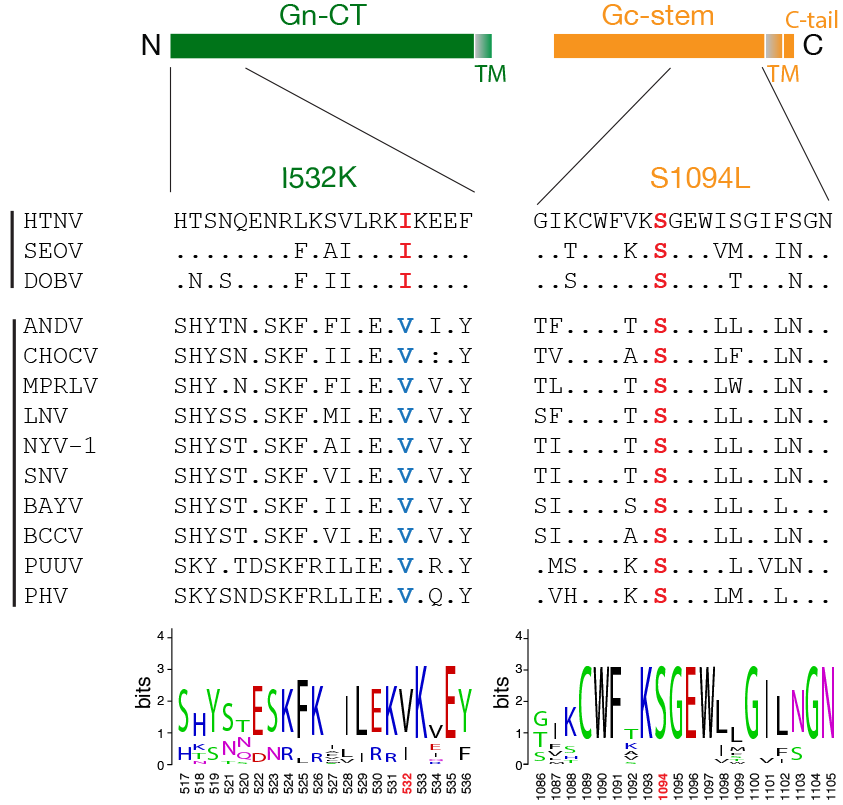
Alignment of Gn/Gc sequences flanking the mutation sites from various hantaviruses. Schematic of cytoplasmic tail of n and stem region of Gc is shown in the top panel. Alignment of amino acid sequences of the N-terminal 20 amino acids of the cytoplasmic tail of Gn and C-terminal 20 acids from 13 species of hantaviruses generated by Clustal Omega along with a WebLogo version (bottom panel) is shown. Abbreviations: Hantaan virus (HTNV), Seoul virus (SEOV), Dobrava-Belgrade virus (DOBV), Andes virus (ANDV), Choclo virus (CHOCV), Maporal virus (MPRLV), Laguna Negra virus (LGNV), New York-1 virus (NYV-1), Sin Nombre virus (SNV), Bayou virus (BAYV), Black Creek Canal virus (BCCV), Puumala virus (PUUV) and Prospect Hill virus (PHV).

How does the I532K mutation enhance cell surface expression of HTNV Gn/Gc? I532 is located in the region that has been shown to bind hantavirus nucleoprotein and RNA (58, 59), just upstream of the dual zinc finger domains in the cytoplasmic tail of the Gn (Gn-CT) protein (**Fig. 7**). Although Golgi retention of many bunyaviruses is mediated by Gn alone (60–65), signals in both hantavirus Gn and Gc seem to contribute to their Golgi localization. Gn proteins of HTNV (39), ANDV or SNV (53, 57) are retained in the ER when expressed alone and need co-expression of Gc for their Golgi transport. On the contrary, Pensiero & Hay (40) reported that HTNV Gn alone can localize to Golgi and the Golgi retention signal is likely located in the N-terminal 20 amino acids of the Gn-CT. The corresponding region of Gc from an orthobunyavirus Uukuniemi virus (UUKV), has also been suggested to be the Golgi retention signal for UUKV Gn/Gc (52). We hypothesize that the I532K mutation relocalizes HTNV Gn/Gc to the cell surface by disrupting its interaction with one or more unknown cellular factors that mediate Golgi retention. The rVSV system described here could be useful for further studies required to characterize this Golgi retention mechanism.

How the Gc (S1094L) mutation increases rVSV-HTNV Gn/Gc infectivity is less clear. It failed to enhance cell surface expression and VSV incorporation of Gn/Gc on its own (**Figs. 3–5**). S1094 is highly conserved across hantaviruses and is located in the membrane-proximal, C-terminal half of the Gc stem (**Fig. 7**). The Gc stem is critical for the formation of the postfusion hairpin conformation (66) and peptides corresponding to its C-terminal half inhibit ANDV infection and membrane fusion (67). Most of the Gc stem, including the S1094 residue was not visualized in the hantavirus Gc crystal structure (68, 69). Alteration in the physical curvature of the membrane by the membrane-proximal region of the VSV glycoprotein stem region has been proposed to enhance VSV budding efficiency (70). However, S1094L alone did not affect budding efficiency (**Fig. 1E, 6A**). We speculate that this mutation might also alter intersubunit interactions and/or the glycoprotein fusogenicity.

Together, our results suggest that the enhancement of cell surface expression of other bunyaviral glycoprotein(s) through incorporation of cognate mutations should enhance the utility of existing single-cycle VSV vectors bearing Old-World hantavirus glycoproteins and facilitate the generation of rVSVs bearing these are other bunyaviral glycoproteins. Moreover, the enhancements in incorporation of Gn/Gc into pseudotyped virus particles and localization at the cell surface in infected cells might elicit a more immunogenic response and pave the way for novel VSV-based bunyaviral vaccines.

## Materials and Methods

### Cells

Human osteosarcoma U2OS and embryonic kidney fibroblast 293T cells obtained from ATCC were cultured in modified McCoy’s 5A media (Thermo Fisher) and high-glucose Dulbecco’s modified Eagle medium (DMEM, Thermo Fisher) supplemented with 10% fetal bovine serum (FBS, Atlanta Biologicals), 1% GlutaMAX (Thermo Fisher), and 1% penicillin-streptomycin (Pen-Strep, Thermo Fisher), respectively. African green monkey kidney Vero cells (from ATCC) were cultured in DMEM supplemented with 2% FBS, 1% GlutaMAX, and 1% Pen Strep. Human umbilical vein endothelial cells (HUVEC, Lonza) were cultured in EGM media supplemented with EGM-SingleQuots (Lonza). All adherent cell lines were maintained in a humidified 37°C, 5% CO_2_ incubator. Freestyle™-293-F suspension-adapted HEK-293 cells (Thermo Fisher) were maintained in GIBCO FreeStyle™ 293 expression medium (Thermo Fisher) using shaker flasks at 115 rpm, 37°C and 8% CO_2_.

### Plasmids

Generation of the plasmid encoding human, codon optimized HTNV Gn/Gc (76-118 strain, GenBank accession number NP_941978.1) in the genome of vesicular stomatitis virus (VSV), carrying an eGFP gene, was described previously (36). HTNV Gn/Gc point mutations (I532K, S1094L, or I532K/S1094L) were cloned into the genomic VSV (described above) and pCAGGS plasmids using standard molecular biology techniques. Human, codon-optimized variable heavy (VH, GenBank accession number FJ751231) and light (VL, GenBank accession number FJ751232) chain sequences of HTNV Gc-specific mAb, 3G1 (37), were synthesized by Epoch Biosciences and cloned into the pMAZ heavy (IgH) and light (IgL) chain vectors (71), respectively. The sequences of all plasmid inserts were confirmed by Sanger sequencing.

### Generation of recombinant and pseudotyped VSVs

Replication-competent, recombinant VSVs (rVSVs) bearing WT or Gn/Gc-mutant HTNV Gn/Gc were generated using a plasmid-based rescue system in 293T cells as described previously (Kleinfelter et al., mBio, Whelan et al., 1995). When required, rescued viruses were propagated on Vero cells, and HTNV Gn/Gc sequences were amplified from viral genomic RNA by RT-PCR and analyzed by Sanger sequencing. Single-cycle VSVΔG pseudotypes, encoding an eGFP reporter, were produced in 293T cells as described previously (Kleinfelter et al., mBio, Whelan et al., 1995). Viral infection was scored by manually enumerating eGFP-expressing cells using an Axio Observer inverted microscope (Zeiss), as described previously (36).

### Production of HTNV Gc-specific mAb 3G1

mAb 3G1 was purified from the supernatants of Freestyle™-293-F suspension cells transiently co-transfected with pMAZ vectors expressing heavy and light chains of 3G1 as described previously (31).

### Detection of HTNV Gn/Gc surface expression by flow cytometry

Human U2OS osteosarcoma cells, seeded in 6-well plates 18-22 h prior to transfection, were transfected with 2 μg of the pCAGGS vectors, expressing nothing or variants of HTNV Gn/Gc, and 0.5 μg of a plasmid expressing eGFP. At 24 h post transfection, cell plates were chilled on ice for 10 min and blocked with chilled 10% Fetal Bovine Serum (FBS) in phosphate buffer saline (PBS) for 30 min at 4°C. Surface HTNV Gc was stained using human anti-HTNV Gc mAb 3G1 (7.3 μg/mL) followed by anti-human AlexaFluor 555 (5 μg/mL, Thermo Fisher) for 1 h at 4°C each. After extensive washing, cells were stained with Live/Dead™ Fixable Violet Dead Cell Stain Kit (Invitrogen), washed again with PBS, and re-suspended in 2% FBS in PBS. Stained cells were passed through a 0.41 μm Nylon Net Filter (Millipore) and analyzed using a LSRII Flow Cytometer and FloJo V.10 software.

### Immunofluorescence microscopy for HTNV Gn/Gc localization

Human U2OS osteosarcoma cells plated on fibronectin-coated glass coverslips were transfected with 500 ng of empty vector or HTNV Gn/Gc expression vectors together with 50 ng of eGFP expressing plasmid as described above. At 24 h post-transfection, cells were fixed with 4% formaldehyde (Sigma) for 5 min and permeabilized with 0.1% Triton X-100 for 10 min at room temperature. After blocking, HTNV Gn and Gc were detected by incubating cells with an anti-HTNV Gn mouse mAb 3D5 (BEI Resources, 1:500 dilution) followed by anti-mouse AlexaFluor-488 antibody, or an anti-HTNV Gc human mAb 3G1 (4.6 μg/mL) followed by anti-human AlexaFluor-555 antibody (Thermo Fisher), respectively. Anti-GM130 (1.25 μg/mL, BD Biosciences) was used to co-stain Golgi apparatus. For surface staining, cells were placed on ice 10 min prior to blocking for 30 min at 4°C and incubated with the above-described HTNV Gn or Gc antibodies on ice before fixing and staining with the secondary antibodies as above. Primary human umbilical vein endothelial cells (HUVECs) were nucleofected with 4.75 μg of empty or HTNV Gn/Gc expressing vectors, together with 50 ng eGFP, using the Amaxa Kit (program A-034, Lonza) before staining for total Gn/Gc expression at 72 h post-nucleofection as described above for U2OS cells. Coverslips were mounted on slides with Prolong containing DAPI (Thermo Fisher) and were imaged using a Zeiss Axio Observer inverted microscope with 40x objective.

### In-cell ELISA for HTNV Gn/Gc expression

293T cells, transfected with empty vector or vectors expressing various HTNV Gn/Gc variants using Lipofectamine-3000 (Invitrogen), were fixed with 4% formaldehyde (Sigma) for 5 min, and permeabilized with 0.1% Triton X-100 for 15 min at room temperature. After blocking with 5% FBS in PBS (1 h at room temperature), total HTNV Gn/Gc expression was detected by incubation with anti-HTNV Gc mAb-3G1 (0.7 μg/mL, 1 h at room temperature) followed by anti-human HRP (Thermo Fisher, 0.045 μg/mL, 1 h at room temperature). ELISA signal was developed using 1-Step™ Ultra TMB-ELISA substrate solution (Thermo Scientific) and measured on a Perkin Elmer Wallac 1420 Victor2™ microplate reader. For measuring cell surface expression of HTNV Gn/Gc, live cells, blocked with 5% FBS in PBS (1 h on ice), were incubated with anti-HTNV Gc mAb-3G1 (0.7 μg/mL, 1 h on ice) before fixing and incubation with the second antibody. No permeabilization step was involved. Absorbance at 450 nm were corrected for background by subtracting the signal from cells transfected with an empty vector.

### ELISA for HTNV Gn/Gc incorporation in VSV particles

To measure HTNV Gn/Gc incorporation into virus particles, we first normalized the ELISA input of single-cycle, pseudotyped vesicular stomatitis viruses (pVSVs) bearing HTNV Gn/Gc variants by immunoblotting (using mouse anti-VSV M mAb 23H12) for the VSV M content. Next, ELISA plates were coated with serial 2-fold dilutions of normalized pVSV particles bearing WT or Gn/Gc mutant HTNV glycoproteins overnight at 4°C. After blocking, HTNV Gc specific was detected using anti-HTNV Gc mAb 3G1 (732 ng/mL) followed by anti-human HRP antibody (44 ng/mL, Thermo Fisher) by incubating for 1 h each at 37°C. ELISA was developed and absorbance at 450 nm was measured as described above.

### Hantavirus Gn/Gc sequence alignment -

Alignment of amino acid sequences of the N-ternimal 20 amino acids of the cytoplasmic tail of Gn and C-terminal 20 acids from 13 species of hantaviruses generated by Clustal Omega. The sequences used for the alignment, along with their GenBank accession numbers, were: Hantaan virus (HTNV) - <u>NP_941978.1</u>; Seoul virus (SEOV) - <u>M34882.1</u>; Dobrava-Belgrade virus (DOBV) - <u>NC_005234.1</u>; Andes virus (ANDV) - <u>NP_604472.1</u>; Choclo virus (CHOCV) - <u>KT983772.1</u>; Maporal virus (MPRLV) - <u>NC_034552.1</u>; Laguna Negra virus (LGNV) - <u>AF005728.1</u>; New York-1 virus (NYV-1) - <u>U36802.1</u>; Sin Nombre virus (SNV) - <u>NP_941974.1</u>; Bayou virus (BAYV) - <u>GQ244521.1</u>; Black Creek Canal virus (BCCV) - <u>L39950.1</u>; Puumala virus (PUUV) - <u>KT885051.1</u> and Prospect Hill virus (PHV) - <u>CAA38922.1</u>. WebLogoes were generated as described earlier (72).

## Acknowledgements

We thank Tyler Krause, Cecelia Harold, and Tanwee Alkutkar for technical support and the Einstein Flow Cytometry Core (supported by NCI center grant P30CA013330). This work is supported by NIH grant AI101436 (to K.C.). K.C. was additionally supported by an Irma T. Hirschl/Monique Weill-Caulier Research Award. M.M.S. was additionally supported by NIH T32 training grant AI070117. The anti-Hantaan virus Gn-specific monoclonal antibody was obtained from the Joel M. Dalrymple - Clarence J. Peters USAMRIID Antibody Collection through BEI Resources, NIAID, NIH: Monoclonal Anti-Hantaan Virus Gn Glycoprotein, Clone 3D5 (produced in vitro), NR-36162.

